# Comparative genomics analysis of *Chryseobacterium* sp. KMC2 reveals metabolic pathways involved in keratinous utilization and natural product biosynthesis

**DOI:** 10.1101/2021.02.23.432615

**Authors:** Dingrong Kang, Saeed Shoaie, Samuel Jacquiod, Søren J. Sørensen, Rodrigo Ledesma-Amaro

## Abstract

Several efforts have been made to valorize keratinous materials, an abundant and renewable resource. Despite these attempts to valorize products generated from keratin hydrolysate, either *via* chemical or microbial conversion, they generally remain with an overall low value. In this study, a promising keratinolytic strain from the genus *Chryseobacterium* (*Chryseobacterium* sp. KMC2) was investigated using comparative genomic tools against publicly available reference genomes to reveal the metabolic potential for biosynthesis of valuable secondary metabolites. Genome and metabolic features of four species were compared, shows different gene numbers but similar functional categories. We successfully mined eleven different secondary metabolite gene clusters of interest from the four genomes, including five common ones shared across all genomes. Among the common metabolites, we identified gene clusters involved in biosynthesis of flexirubin-type pigment, microviridin, and siderophore, all showing remarkable conservation across the four genomes. Unique secondary metabolite gene clusters were also discovered, for example, ladderane from *Chryseobacterium* sp. KMC2. Additionally, this study provides a more comprehensive understanding of the potential metabolic pathways of keratin utilization in *Chryseobacterium* sp. KMC2, with the involvement of amino acid metabolism, TCA cycle, glycolysis/gluconeogenesis, propanoate metabolism, and sulfate reduction. This work uncovers the biosynthesis of secondary metabolite gene clusters from four keratinolytic *Chryseobacterium* spp. and shades lights on the keratinolytic potential of *Chryseobacterium* sp. KMC2 from a genome-mining perspective, providing alternatives to valorize keratinous materials into high-value natural products.

**Importance:** Keratin is an abundant and renewable resource from slaughterhouses or the poultry industry. Low-value products such as animal feed and fertilizer were generated from these feedstocks based on conventional processing like chemical conversion. In fact, microorganisms possess the potential to synthesize valuable natural products. In this work, we explored the metabolic potential of *Chryseobacterium* sp. KMC2, which was isolated with efficient keratinolytic capacity from a previous study. Comparative genomics analysis displayed similar functional categories against three publicly available reference genomes of keratin-degrading *Chryseobacterium* spp.. Eleven different secondary metabolite gene clusters of interest were mined among four genomes, including five common and unique ones. Furthermore, we provide a more comprehensive understanding of metabolic pathways on keratin utilization in *Chryseobacterium* sp. KMC2, with the involvement of amino acid assimilation and sulfate reduction. These findings contribute to expanding the application of *Chryseobacterium* sp. KMC2 on the valorization of keratinous materials.

## Introduction

Keratin is the most abundant proteins in epithelial cells, constituting the bulk of epidermal appendages such as hair and feather (1, 2). Keratinous materials represent an abundant protein source, particularly originating from the commercial slaughterhouses or poultry farms (3). They contain peptides and amino acids, which are renewable natural resources with great potential in sustainable development (4). However, keratin is an insoluble protein with highly cross-linked disulfide bonds giving it a tough and recalcitrant structure (5). Many attempts have been made to hydrolysis keratinous materials in terms of physicochemical treatment, enzymatic hydrolysis, and microbial conversion (6, 7). The hydrolysis products of keratinous materials have been used for animal feed (8) and fertilizer (6, 9) based on conventional processing.

Microorganisms represent one of the most important natural sources, which have the potential to generate bioactive compounds such as antibiotics, biofuels, and natural pigments derived from cellular metabolites (10, 11). For example, *Yarrowia lipolytica* has been used to convert different renewable feedstocks to high-value metabolites (12). Similarly, *Escherichia coli* has become one of the best cell reactors to produce alcohols, organic acids, biodiesel, even hydrogen by utilizing renewable resources (13). Other bacteria such as *Bacillus subtilis* (14), *Caldicellulosiruptor bescii* (15), *Corynebacterium glutamicum* (16), and *Ruminococcaceae* bacterium (17) were identified and evaluated with the capacity to generate different products by converting renewable carbon sources. Notably, some microorganisms were reported to degrade keratinous waste effectively (18). Exploring keratinolytic potential of these microbes to generate high value-added products is an important step to recycle and valorize keratinous materials.

Molecular mechanisms of microbial keratin degradation are still not fully understood, while genome sequencing offers possibilities to reveal the metabolic potential behind efficient microbial degradation (19). Novel keratinolytic enzymes were identified from the genome of *Bacillus pumilus* 8A6, an efficient keratin degrader (20). Furthermore, going beyond the degradation reaction itself, genomes can also be minded for valuable accessory functions of interest, adding more values to the microbial conversion processes. For instance, gene clusters and biosynthesis pathways of secondary metabolites could be disclosed from genomes *via* adequate analysis tools (21, 22). A total of 104 putative biosynthetic gene clusters (BGCs) for secondary metabolites were predicted from nine *Ktedonobacteria* genomes (23). Secondary metabolites were identified and linked to gene clusters based on the comparison and mining of six genomes belonging to diverse *Aspergillus* species, successfully fueling industrial biotechnology initiatives and medical research (24). Therefore, using the genomes of keratinolytic microbial species in a similar way would represent a promising approach to discover biosynthetic gene clusters of secondary metabolites of interest, excavating the full application potential of these microbes.

Recently, several studies based on different environments have revealed the remarkable potential of representative taxa from the *Chryseobacterium* genus for keratin degradation using isolation, activity tests and genome sequencing (25, 26). In this study, a novel strain *Chryseobacterium* sp. KMC2 was previously obtained from an enrichment procedure, displaying a potent capacity of keratin degradation (27). The genome of *Chryseobacterium* sp. KMC2 was analyzed and compared with publicly available genomes of other keratinolytic *Chryseobacterium* spp. to clarify the genomic basis of keratin degradation, and to unravel hidden biosynthetic gene clusters of interest. Subsequently, the metabolic pathways associated with keratin degradation were constructed, providing deeper insight into the yet obscure keratinolytic processes. This work reveals the keratinolytic potential of *Chryseobacterium* spp. and mined potential accessory gene clusters of secondary metabolites, which could i) contribute to optimizing the processes of keratin degradation and ii) broaden the perspective to generate added-value products from keratin hydrolysate.

## Results

### Genome feature comparison of four keratinolytic *Chryseobacterium* strains

*Chryseobacterium* sp. KMC2 originated from a river-bank soil sample, and displayed a potent degradation ability toward milled pig bristle and hooves (27, 28). The genome of *Chryseobacterium* sp. KMC2 was sequenced and compared to three reference genomes of *Chryseobacterium* spp. Including: i) *Chryseobacterium camelliae* Dolsongi-HT, isolated from green tea leaves (29); ii) *Chryseobacterium gallinarum* strain DSM 27622, isolated from chicken (30); and iii) *Chryseobacterium* sp. strain P1-3 isolated from poultry waste (31), which all display keratinolytic capacity (Table 1). *Chryseobacterium* sp. KMC2 showed distinct genome feature from the other known keratinolytic strains. The genome size of *Chryseobacterium* sp. KMC2 is 5.28 Mbp, larger than the other three genomes, which ranged from 4.38 Mbp to 4.63 Mbp. A total of 4,773 genes were predicted from *Chryseobacterium* sp. KMC2 genome, and about 4,000 genes were annotated from the other three genomes. Besides, the GC content ranges from 36.33% to 41.80% in *Chryseobacterium* spp. genomes. Furthermore, the whole-genome phylogenetic tree was constructed with other eight publically available *Chryseobacterium* spp. genomes (Fig. 1a), showing the highly similarity among *Chryseobacterium gallinarum* strain DSM 27622 and *Chryseobacterium* sp. strain P1-3. Notably, *Chryseobacterium* sp. KMC2 and *Chryseobacterium camelliae* Dolsongi-HT have closer evolutionary relationship with other *Chryseobacterium* species. Although the keratinolytic capacity within these eight strains is still not clear, more keratinolytic species are expected to be found in this genus according to the phylogenic tree analysis. Four strains among this genus have documented with actual keratin degradation capability (29-31), while *Chryseobacterium* sp. KMC2 may possess different keratinolytic potential encoded in its larger genome compared to three other *Chryseobacterium* strains.

**Table 1.** Features comparison of four *Chryseobacterium* spp. genomes.

|  | *Chryseobacterium* | *Chryseobacterium camelliae* | *Chryseobacterium gallinarum* | *Chryseobacterium* |
| --- | --- | --- | --- | --- |
| Parameters | sp. KMC2 | Dolsongi-HT1 | strain DSM 27622 | sp. P1-3 |
| **Total length (bp)** | 5.276.159 | 4.376.354 | 4.633.632 | 4.628.764 |
| **Contigs** | 63 | 1 | 1 | 45 |
| **N50 (bp)** | 231.784 | 4.376.354 | 4.633.632 | 342.512 |
| **GC content (%)** | 36.33 | 41.80 | 37.30 | 37.02 |
| **Gene** | 4.773 | 3.999 | 4.182 | 4.087 |
| **CDS** | 4.692 | 3.881 | 4.033 | 3.119 |

**Fig. 1.**
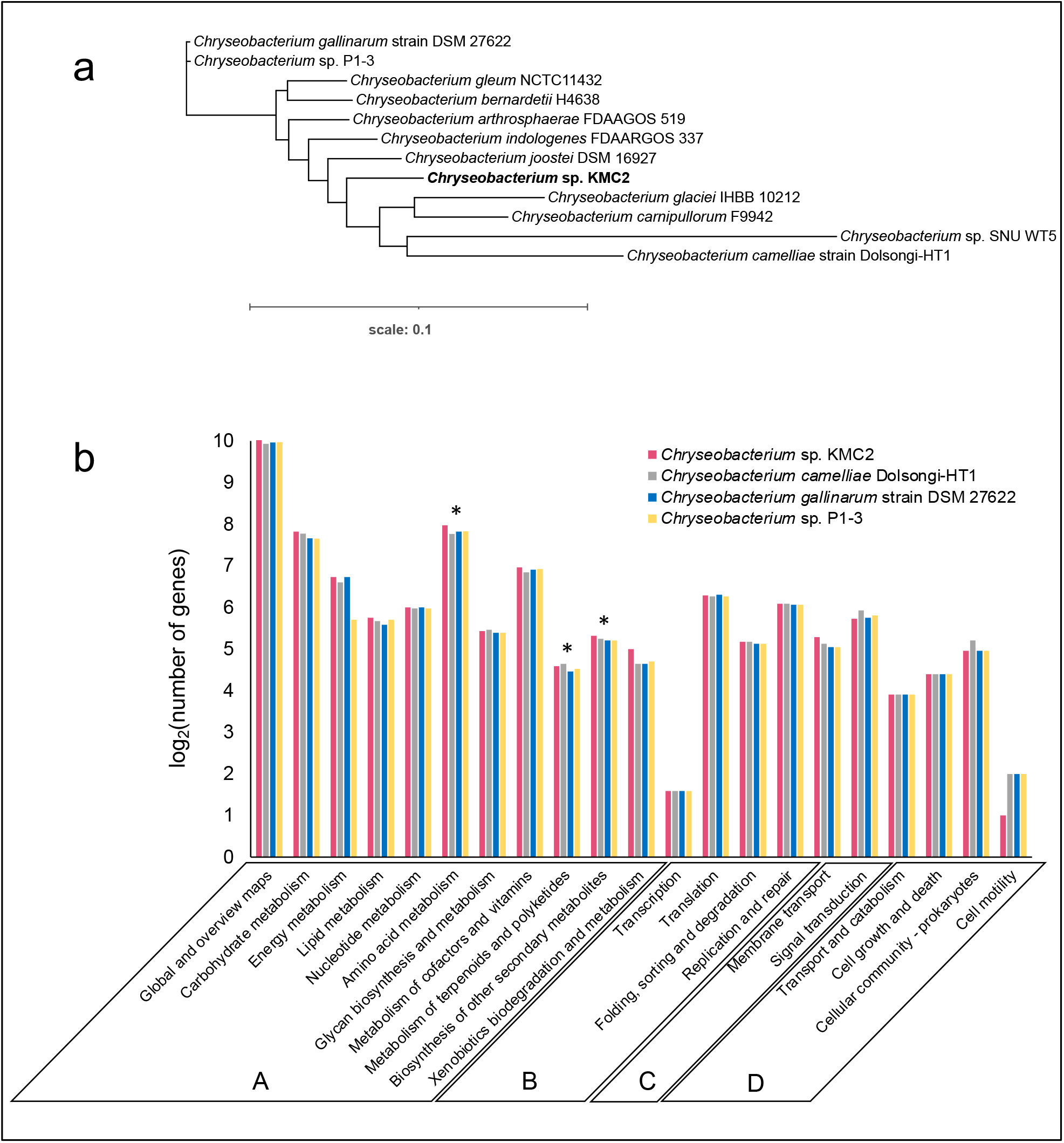
Analysis of the *Chryseobacterium* spp. genomes. (a). The whole-genome sequence-based phylogenetic tree of *Chryseobacterium* spp., constructed by REALPHY 1.12 (62), based on the merge reference alignments of all genomes. Branch length represents divergence. (b). Comparison of KEGG function classification amongst four *Chryseobacterium* spp. genomes. Functional categories: Metabolism (A), Genetic information processing (B), Environmental information processing (C), and Cellular processes (D). The stars show the sub-categories: Amino acid metabolism, metabolism of terpenoids and polyketides, and biosynthesis of other secondary metabolites.

### Metabolic potential comparison of four keratinolytic *Chryseobacterium* genomes

About 40% of the genes from the four genomes were classified into various functional categories based on the KEGG database. The vast majority of annotated genes belonged to metabolism, genetic information processing, environmental information processing, and cellular processes (Fig. 1b). The functional categories of the genomes were overall highly similar, with ∼85% of annotated genes assigned to “metabolism” (category A) which included ∼1.000 genes into the sub-category “global and overview maps”. Additionally, about 8% and 4% annotated genes from each genome were assigned to “genetic information processing” (category B) and “environmental information processing” (category C), respectively. The remaining annotated genes belonged to “cellular processes” (category D), which occupied 3% of the annotated genomes approximately.

Remarkably, each genome had more than 200 genes assigned into the “amino acid metabolism” sub-category. Keratin is mainly composed of amino acids (3), which is ultimately the operational carbon nutrient source exploited for microbial growth. Numerous amino acid metabolism-related enzymes were annotated, revealing the genetic potential of these *Chryseobacterium* strains for using keratin materials as energy sources. Of particular interest, several biosynthesis genes of secondary metabolites were detected from the genomes, of which more than 20 genes were annotated as “metabolism of terpenoids and polyketides” and around 40 genes were annotated as “biosynthesis of other secondary metabolites” sub-category (Fig. 1b). Terpenoids are a group of natural products with diverse commercial applications, which have been produced from microbial cell factories (32). Many polyketides are considered as significant natural products with broad applications in the agriculture and pharmaceutical industry (33). The metabolic pathways related to polyketides biosynthesis production are well understood in some microorganisms like *Streptomyces* which play a crucial role in industrial bioproduction (34). This result indicates that these *Chryseobacterium* strains could have the potential to synthesize high-value secondary metabolites such as terpenoids and polyketides from keratinous materials.

### Mining and comparing secondary metabolite gene clusters

Genome mining is an effective approach to discover new natural products from microorganisms based on “signature genes” detection or searching for specific patterns in gene sequences (35). To explore the potential of producing high value chemicals from these four *Chryseobacterium*, secondary metabolite gene clusters were predicted by using antiSMASH 5.0 mining pipeline (Fig. 2). In total, eleven different secondary metabolite gene clusters were identified. *Chryseobacterium* sp. KMC2 possesses the largest number (15), while *Chryseobacterium camelliae* Dolsongi-HT1 has the fewest (8). Ten gene clusters were predicted from the other two strains. Five clusters are present in the four genomes, which are flexirubin-type pigment (resorcinol and arylpolyene), microviridin, lanthipeptide, NRPS-like, and siderophore. Remarkably, the flexirubin-type pigment is a typical metabolite produced from *Flavobacterium* (36). Several species from *Chryseobacterium* were previously designated and known as *Flavobacterium* owes to similar characteristics including the presence of yellow pigments (37). Flexirubin-type pigment was isolated and characterized from *Chryseobacterium* sp. UTM-3T (38). In addition, *Chryseobacterium* sp. KMC2 owns a unique gene cluster to produce ladderane. Another unique natural product is beta-lactone from *Chryseobacterium camelliae* Dolsongi-HT1. Ladderanes are hydrocarbon chains which were regarded as membrane lipid components produced by anammox (anaerobic ammonia-oxidizing) bacteria uniquely, but the production is not affordable due to their extremely low growth (39, 40). These results demonstrate that various secondary metabolite gene clusters including both expected and unusual candidates were discovered from *Chryseobacterium* genomes, which could turn into novel natural product sources.

**Fig. 2.**
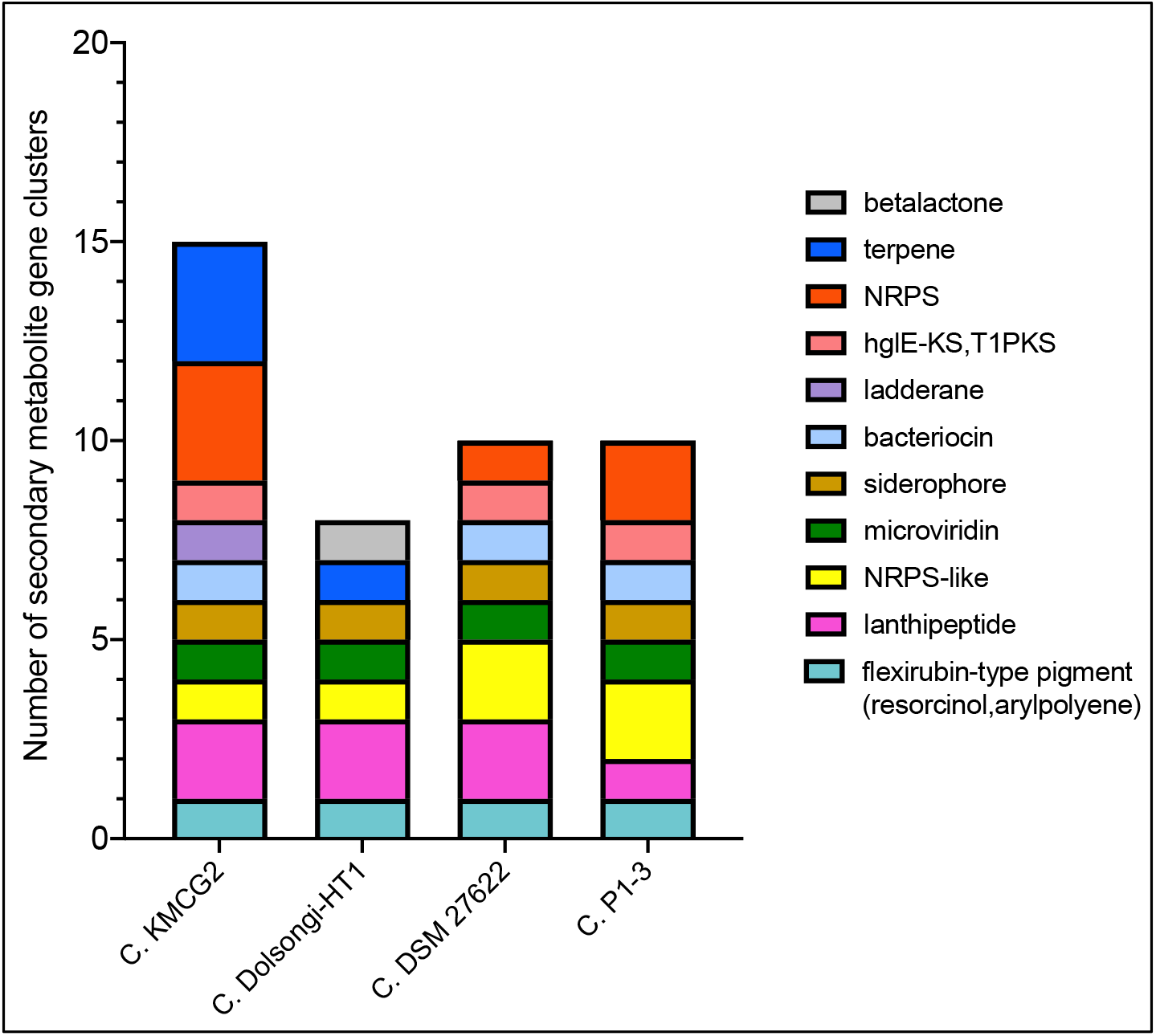
Composition of secondary metabolite gene clusters from four *Chryseobacterium* spp. genomes.

### Synteny analysis and features of secondary metabolite gene clusters

Comparative genomics can reveal unique cluster and distribution patterns of secondary metabolites in species (41). Five secondary metabolite gene clusters, predicted to be present in the four genomes, were selected to explore the evolutionary relationship among four *Chryseobacterium* strains. Three of them including flexirubin-type pigment, microviridin, and siderophore display a conserved gene cluster structure from synteny analysis (Fig. 3, 4, and 5), while the other two showed no evident synteny relation (Fig. S1 and S2).

**Fig. 3.**
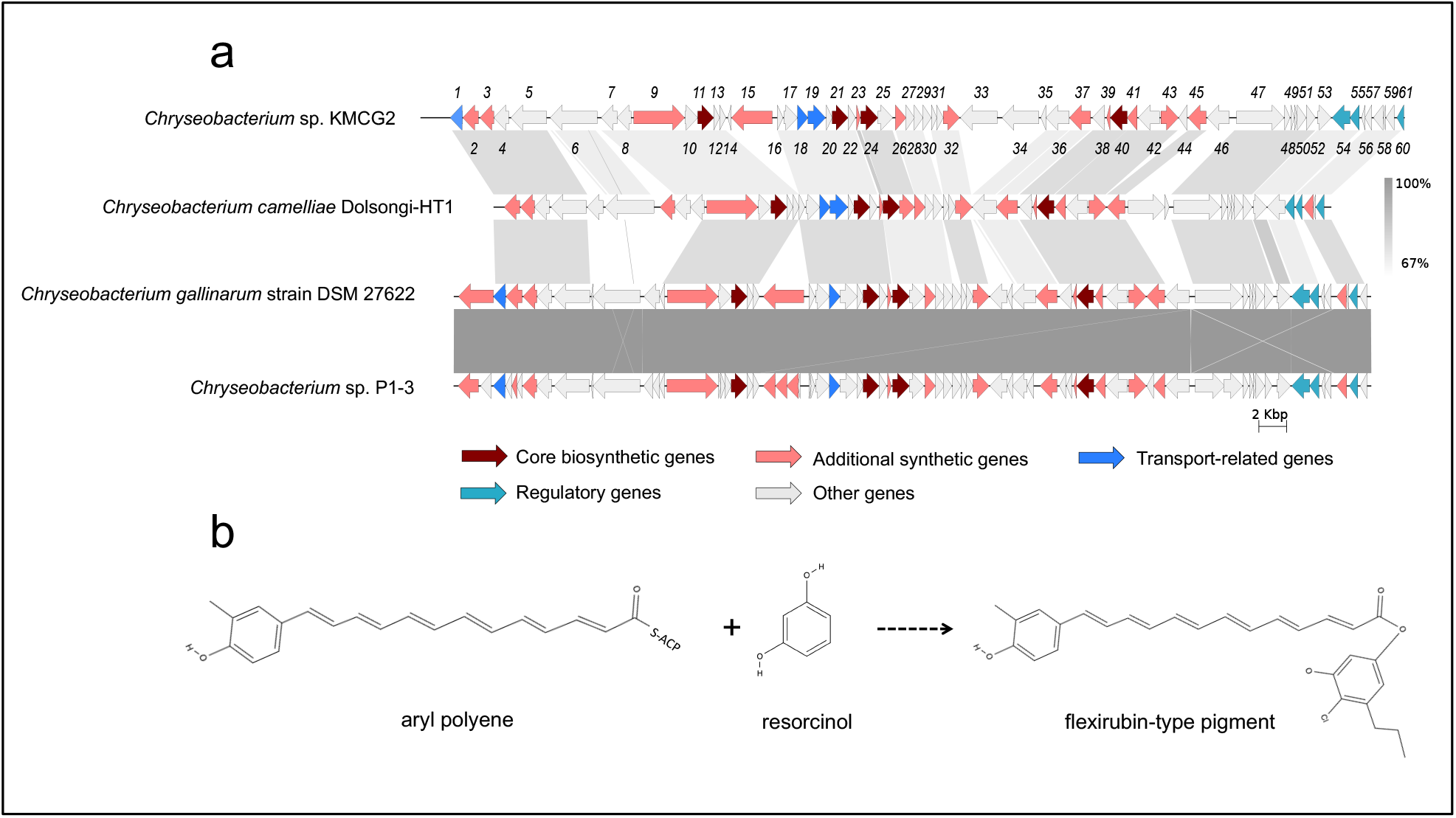
Flexirubin-type pigment gene cluster from four *Chryseobacterium* spp. genomes. (a). Synteny analysis and features of flexirubin-type pigment gene cluster in *Chryseobacterium* spp. genomes. (b). The proposed biosynthetic reaction of flexirubin-type pigment. The detailed description of each gene can be found in Table S1.

**Fig. 4.**
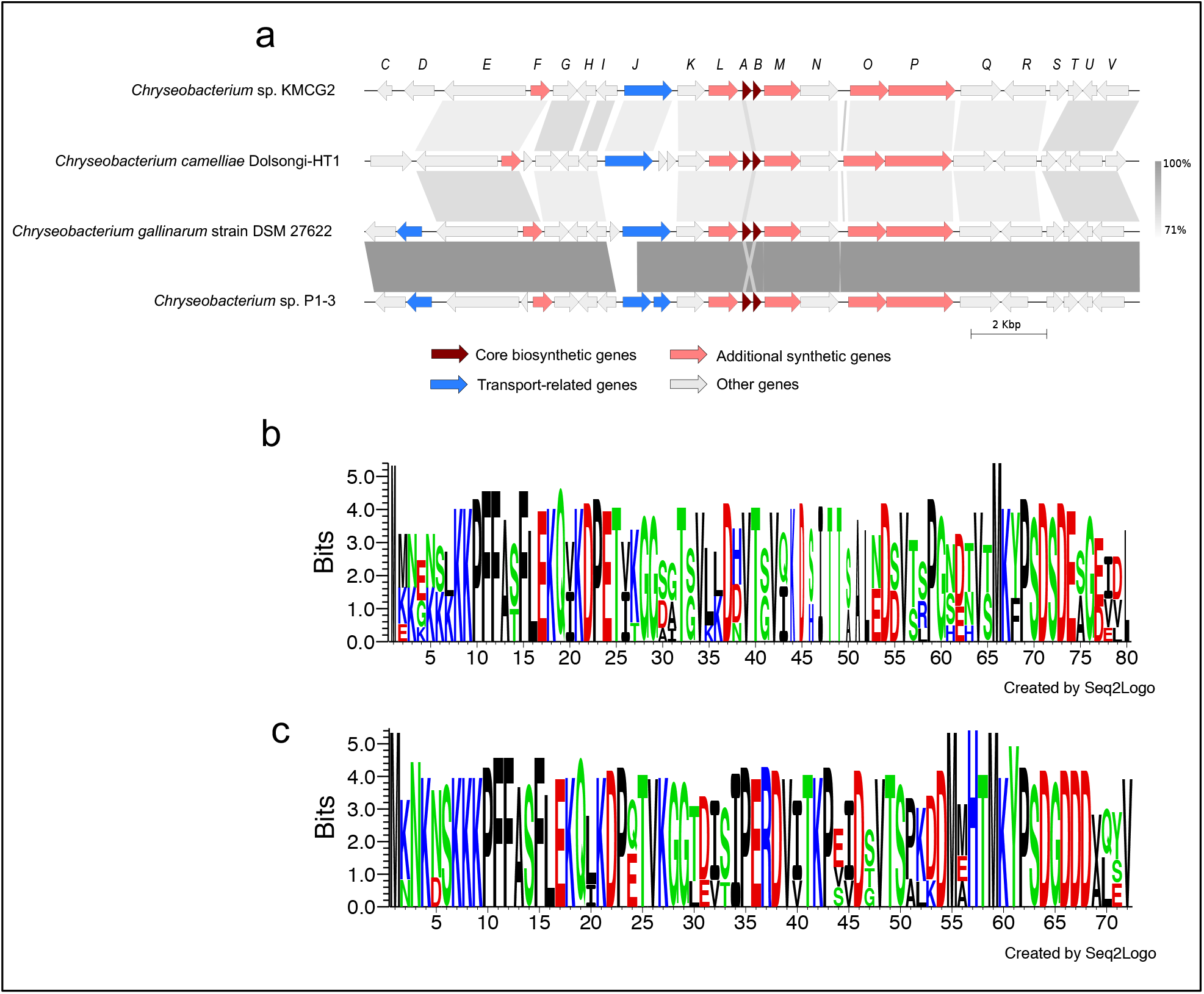
Microviridin gene cluster from four *Chryseobacterium* spp. genomes. (a). Synteny analysis and features of microviridin gene cluster in *Chryseobacterium* spp. genomes. (b). Amino acid sequence comparison of mdnA from four *Chryseobacterium* spp. genomes. (c). Amino acid sequence comparison of mdnB from four *Chryseobacterium* spp. genomes. The detailed description of each gene can be found in Table S2.

**Fig. 5.**
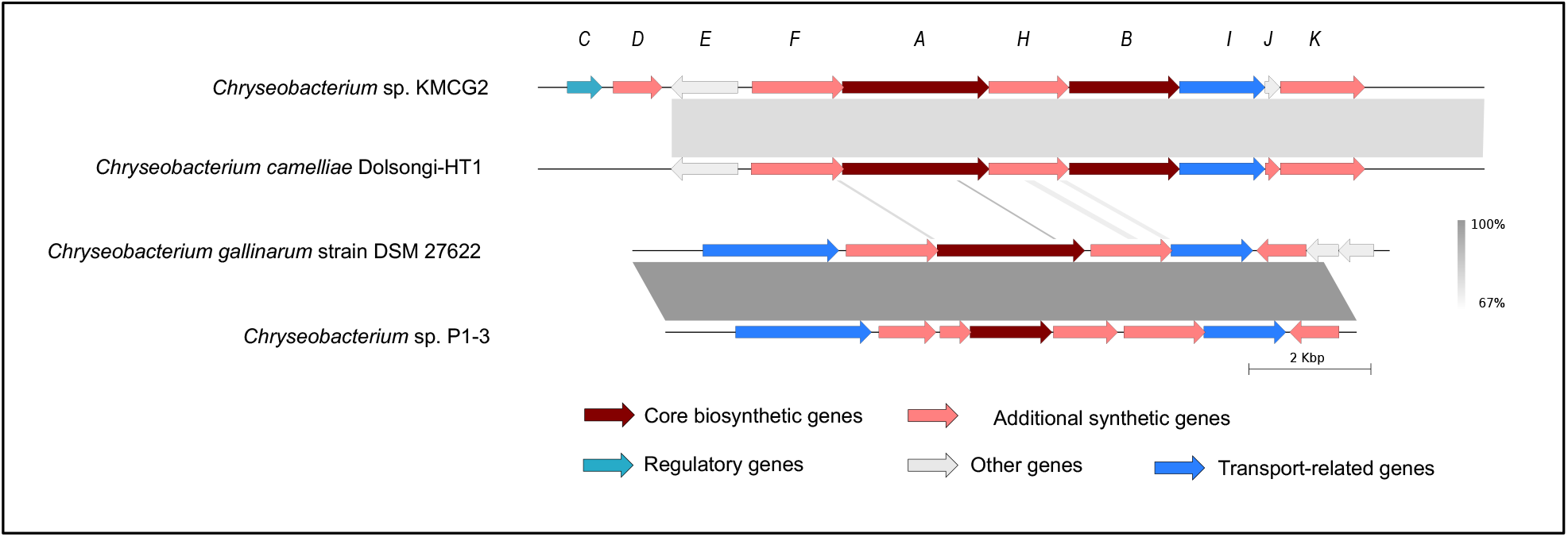
Siderophore gene cluster from four *Chryseobacterium* spp. genomes. The detailed description of each gene can be found in Table S3.

### Flexirubin-type pigment

Natural pigments have increasing applications in food, pharmaceutical, and textile industries, owing to their advantages such as non-toxic, biodegradable, and low allergenic potential compared to synthetic pigments (42). In particular, flexirubin-type pigment has a potential antimicrobial and anti-tumoral activities (43). Biosynthesis gene clusters of flexirubin-type pigment are conserved across the four tested genomes, especially within *Chryseobacterium gallinarum* strain DSM 27622 and *Chryseobacterium* sp. strain P1-3 (Fig. 3a). A total of 61 biosynthesis-related genes of flexirubin-type pigment were predicted from *Chryseobacterium* sp. KMC2, including four core biosynthesis genes. One of the core biosynthesis genes was annotated as 3-oxoacyl-(acyl carrier protein) synthase III (*Flex11*), and the other three were annotated as Beta-ketoacyl synthases (*Flex21, Flex24*, and *Flex40*). Besides, transport-related genes and regulatory genes were predicted from the gene cluster. A previous study identified the molecular structure of flexirubin-type pigment isolated from a *Chryseobacterium* sp. UTM-3T (38). According to the products from core biosynthesis genes and their molecular structures, a proposition of biosynthesis pathway was established (Fig. 3b), where flexirubin-type pigment is generated from resorcinol and arylpolyene. Further transcriptomics and metabolomics analysis would be required to confirm the validity of this potential pathway discovery.

### Microviridin

Microviridins represent a group of ribosomally synthesized peptides under post-translational modifications, which have been mainly isolated from cyanobacteria and present potent serine-type protease inhibitory activities (44, 45). These properties could make microviridin serve as the natural antimicrobial agents for developing potential drugs. Biosynthesis gene clusters of microviridin from four *Chryseobacterium* genomes show a highly conserved structure with a similarity greater than 71% from most gene synteny analysis (Fig. 4a). There are 23 biosynthesis genes of microviridin that were predicted from *Chryseobacterium* sp. KMC2. Two core biosynthetic genes (A and B) were identified from genomes and transport-related genes were also been discovered. Besides, amino acid sequences of mvdA and mvdB were aligned, showing that multiple motifs from mvdA and mvdB are conserved (Fig. 4bc). Interestingly, many keratinases were reported to be classified as serine proteases, acting on the molecular structure of keratin (46). This suggests that microviridins may regulate keratinolytic activity, further characterizing and manipulating the microviridin synthetic pathway could contribute to improving the keratin degradation efficiency.

### Siderophore

Siderophores are ferric ion-specific chelators to scavenge iron from the extracellular environment, which play important roles in virulence and oxidative stress tolerance in microorganisms (47). It has been designed as a Trojan horse antibiotic to enter and kill pathogenic bacteria (48), and also has shown the potential to decrease the growth of cancerous cells (49). Biosynthesis gene cluster of siderophore shows a high synteny conservation among *Chryseobacterium* sp. KMC2 and *Chryseobacterium camelliae* Dolsongi-HT1, *Chryseobacterium gallinarum* strain DSM 27622 and *Chryseobacterium* sp. Strain P1-3, respectively (Fig. 5). A total of ten genes were predicted from siderophore biosynthesis cluster of *Chryseobacterium* sp. KMC2, and eight genes from the other three *Chryseobacterium* strains separately. Functional description of each gene related to siderophore biosynthesis in *Chryseobacterium* sp. KMC2 shows two core biosynthesis genes, and includes one regulatory gene and one transport-related gene. This further suggests that those siderophore are potentially fully functional molecular features that can be regulated on-demand and exported outside the cell when needed.

### Metabolic pathways of keratin utilization in *Chryseobacterium* sp. KMC2 genome

The main metabolic pathways related to keratin utilization in *Chryseobacterium* sp. KMC2 genome were investigated. These pathways included amino acid metabolism, TCA cycle, glycolysis/gluconeogenesis, propanoate metabolism, and sulfate reduction (Fig. 6). A previous study suggested that abundant amino acids are released during microbial degradation and used as nutrient sources, such as leucine and aspartate (28). The metabolic pathways of amino acid utilization were mapped from the genome of *Chryseobacterium* sp. KMC2. Most of the amino acids are converted into intermediates of the TCA cycle. For instance, arginine can be converted to succinate, then enter to TCA cycle after a multiple-steps enzyme reaction. Aspartate, tyrosine, phenylalanine, and glutamate could serve as the substrates to generate fumarate, thus being part of the TCA cycle. Besides, isoleucine turns into the substrates of 2-methyl-acetoacetyl-CoA after several enzymatic steps, which is then converted into acetyl-CoA and propanoyl-CoA *via* acetyl-CoA C-acyltransferase. Acetyl-CoA is an important intermediate, which can entry to the TCA cycle *via* citrate synthase (50). It is also the precursors of fatty acid and polyketides biosynthesis (51). Propanoyl-CoA serves as the critical substrate within propanoate metabolism and can also be used to make lipids (52, 53). On the other hand, methionine can be converted to 2-oxobutanoate, which is also an intermediate of propanoate metabolism. Subsequently, the methylmalonyl-CoA generated in propanoate metabolism enters into the TCA cycle *via* succinyl-CoA. Besides, the key enzymes of glycolysis/gluconeogenesis were found, indicating the potential to produce essential biomass components based on oxaloacetate from TCA cycle. Evidence indicates that a source of redox is needed for complete keratin degradation with keratinases (54, 55). Several metabolites such as sulfite were revealed to be associated with efficient keratin degradation (56). Therefore, here the complete metabolic pathway of sulfate reduction was mapped in the genome, which shows that the potential to create a redox environment needed for keratinases is indeed present. Following the development of sequencing technologies, increasing genomes of keratinolytic species have been unveiled, which provide a genomic perspective to reveal the molecular keratinolytic mechanisms. For instance, metabolic pathways related to keratin degradation such as enzymolysis and reduction of disulfide bonds were clarified through uncovering the genetic basis of microbial genomes (57). The complex keratinolytic processes of *Streptomyces* sp. included protease secretion, iron uptake, spore formation, and resuscitation were recently revealed from a genome view (58). Our results are in line with the notion that a redox environment is indeed required for efficient keratinolytic activity to occur. It is expected that the integrated metabolic pathways associated with keratinolytic processes will be deciphered along with more genomes sequencing and biochemical studies of relevant metabolic pathways.

**Fig. 6.**
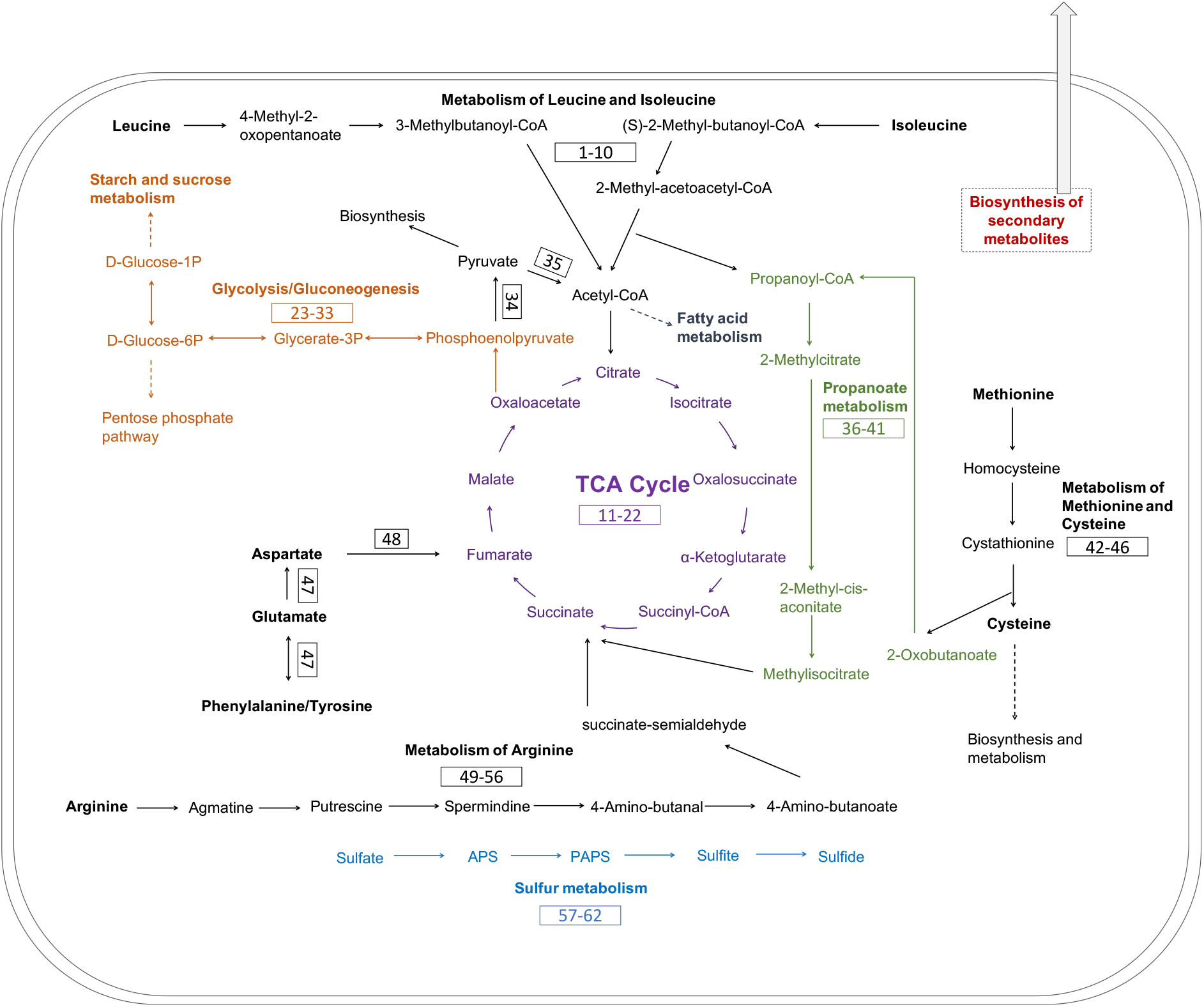
Metabolic pathways reconstruction of keratin utilization in *Chryseobacterium* sp. KMC2 genome. It includes amino acid metabolism, TCA cycle, glycolysis/gluconeogenesis propanoate metabolism, and sulfur metabolism. The number in the box represents the gene related to the metabolic pathway. The detailed description of each gene can be found in Table S7.

## Conclusion

In this work, the genomes from four *Chryseobacterium* spp. with keratinolytic activity were analyzed. Common and unique secondary metabolite gene clusters were mined from *Chryseobacterium* spp. genomes, suggesting the potential to generate high value metabolites using keratin-rich wastes as the nutrient sources. Therefore, the use of these microorganisms could be an alternative way to valorize keratinous materials through microbial conversion. Furthermore, the metabolic pathways of keratin degradation from *Chryseobacterium* sp. KMC2 was studied from a genomic viewpoint. Nevertheless, there are still unknowns to link both metabolic pathways of keratinous utilization and the natural products biosynthesis. Understanding these connected pathways and their regulation will contribute to developing synthetic biology approaches to boost high value-added products from microbial keratin degradation.

## Materials and Methods

### DNA preparation

*Chryseobacterium* sp. KMC2 was isolated and identified from a keratinolytic microbial consortium enriched from a soil sample (19, 27, 28). The keratinolytic capacity of *Chryseobacterium* sp. KMC2 was evaluated as the keystone strain among the microbial consortium, which was able to degrade keratin materials efficiently (27). *Chryseobacterium* sp. KMC2 was inoculated to LB medium, and cultured overnight (200 rpm, 30 °C). Two milliliters of the suspension were centrifuged and collected to prepare the DNA extraction, performed by using by FAST Soil DNA Kit (MP Biomedicals, United States) according to the manufacturer’s instructions.

### Genome sequencing, assembling, and functional annotation

The genome sequencing was performed by an Illumina Miseq instrument at the University of Copenhagen by using DNA Library Preparation Kits v2 (2 × 250 bp), according to the manufacturer’s instructions. Raw reads were treated and assembled to contigs on CLC Genomic Workbench 8.5.1. The obtained contigs were validated using QUAST 4.5 (59). Genes were predicted from the contigs and further annotated with Prokka v1.14.5 (60). Predicted genes from contigs were submitted to eggNOG 5.0 database to obtain an integrated functional annotation and classification (61).

### Whole-genome phylogenetic analysis

To determine the phylogenetic origin of *Chryseobacterium* sp. KMC2 in the *Chryseobacterium* genus, the whole-genome sequences of 11 publicly available *Chryseobacterium* spp. were downloaded from NCBI database to construct a phylogenetic tree. The whole-genome sequence-based phylogenetic tree was inferred by using an online pipeline: The Reference sequence Alignment based Phylogeny builder (REALPHY 1.12) (62), based on the merge reference alignments. Then the visualization of the phylogenetic tree was generated by iTOL v5 (63).

### Secondary metabolite gene cluster detection and annotation

Assembled contigs of four *Chryseobacterium* spp. were uploaded to antiSMASH 5.0 secondary metabolite genome mining web platform (21). Predicted secondary metabolites gene clusters from *Chryseobacterium* sp. KMC2 were compared with other keratinolytic *Chryseobacterium* strains. Gene annotation of each cluster from *Chryseobacterium* sp. KMC2 was performed by Prokka v1.14.5 (60) and BLASTP with the NCBI database. The best match sequencing ID was recorded for the annotated genes. Synteny and features of conservative secondary metabolite gene clusters were analyzed by using Easyfig 2.2.2, showing the similarity of gene sequences (64). Feature comparison of amino acid sequences and motifs from core synthetic genes were analyzed by using Clustal Omega (65) to get the multiple sequence alignment and using Seq2logo to generate sequence logo (66).

### Metabolic networks construction

The genomes of *Chryseobacterium* sp. KMC2 and other three *Chryseobacterium* spp. were submitted to GhostKOALA (67) to obtain the KO number for each gene, then genes were assigned to different metabolic pathways and functional categories. Following the metabolic networks construction of *Chryseobacterium* sp. KMC2 was achieved through mapping the annotated enzyme genes to KEGG (68) reference pathway and Biocyc database (69) manually.

## Data availability

Reference genomes were downloaded from the NCBI database: *Chryseobacterium camelliae* strain Dolsongi-HT1 (GenBank: GCA_002770595.1), *Chryseobacterium gallinarum* strain DSM 27622 (GenBank: GCA_001021975.1), *Chryseobacterium* sp. P1-3 (GenBank: GCA_000738495.1), *Chryseobacterium gleum* NCTC11432 (GenBank: GCA_900636535.1), *Chryseobacterium bernardetii* H4638 (GenBank: GCA_003815955.1), *Chryseobacterium* arthrosphaerae FDAAGOS 519 (GenBank: GCA_003812705.1), *Chryseobacterium indologenes* FDAARGOS 337 (GenBank: GCA_002208925.2), *Chryseobacterium joostei* DSM 16927 (GenBank: GCA_003815775.1), *Chryseobacterium glaciei* IHBB 10212 (GenBank: GCA_001648155.1), *Chryseobacterium carnipullorum* F9942 (GenBank: GCA_003815855.1), and *Chryseobacterium* sp. SNU WT5 (GenBank: GCA_007362475.1). Raw sequencing data were deposited in the Sequence Read Archive (SRA) database under the BioProject number PRJNA686768 with an accession number SRR13278108. The assembled genome sequence of *Chryseobacterium* sp. KMC2 has been deposited at DDBJ/ENA/GenBank under the accession JAESIT000000000.

## Acknowledgements

This research was funded by the Danish Innovation Fund (Grant Number 1308-00015B, Keratin2Protein) and also under the support of the Chinese Scholarship Council (CSC) Scholarship Program.

## Notes

### Competing Interest Statement

The authors have declared no competing interest.

### Summary of Updates

Supplemental files updated

