## Supplemental Material for "Comparative genomics analysis of *Chryseobacterium* sp. KMC2 reveals metabolic pathways involved in keratinous utilization and natural product biosynthesis"

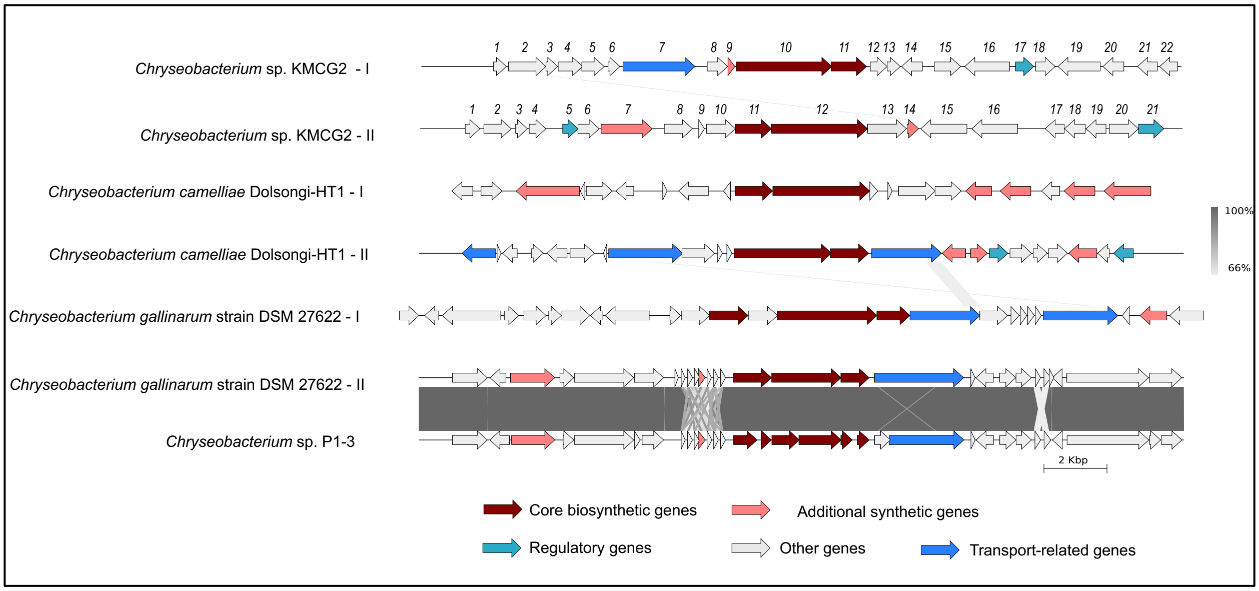


**Fig. S1.** Lanthipeptide gene cluster from four *Chryseobacterium* spp. genomes (Two gene clusters were predicted to synthesize the same secondary metabolite showing as “I” and “II”, respectively). The detailed description of each gene can be found in Table S4&5.

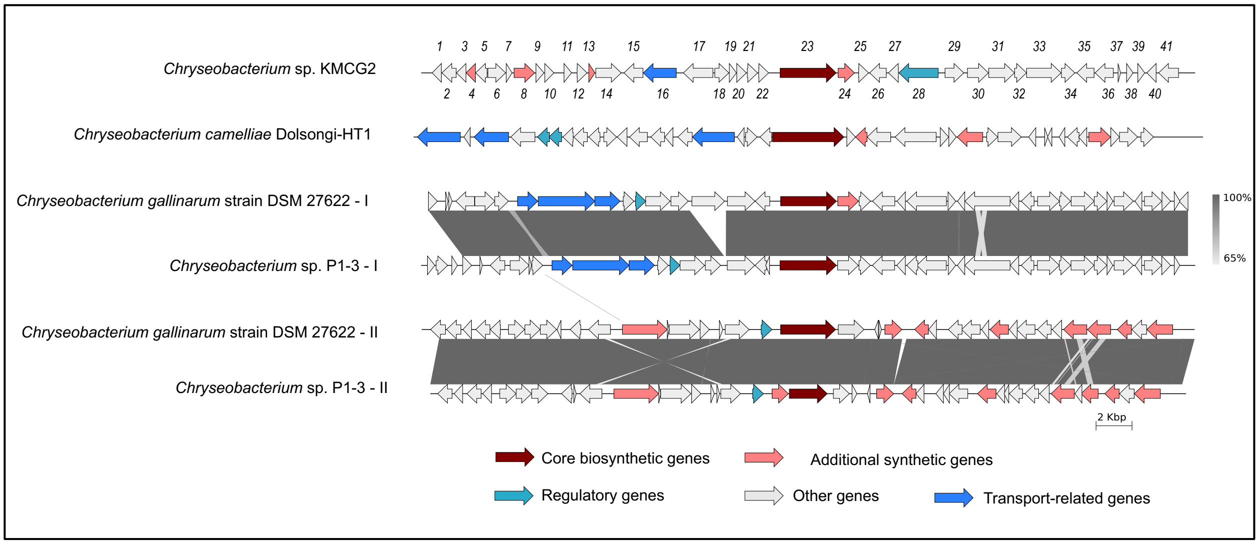


**Fig. S2.** NRPS-like gene cluster from four *Chryseobacterium* spp. genomes (Two gene clusters were predicted to synthesize the same secondary metabolite showing as "I" and "II", respectively). The detailed description of each gene can be found in Table S6.

**Table S1.** The description of flexirubin-type pigment biosynthesis related gene cluster from *Chryseobacterium* sp. KMC2

| Genes | Length (bp) | Functional annotation (Sequence ID from NCBI BLASTP) | Reference in genome |
|---|---|---|---|
| *flex1* | 876 | Polysaccharide export protein | KOACACPH_03512 |
| *flex2* | 1119 | Decaprenyl-phosphate N-acetylglucosaminephosphotransferase | KOACACPH_03513 |
| *flex3* | 1002 | Undecaprenyl-phosphate 4-deoxy-4-formamido-L-arabinose transferase | KOACACPH_03514 |
| *flex4* | 1071 | Formimidoylglutamase | KOACACPH_03515 |
| *flex5* | 2580 | DNA topoisomerase 1 | KOACACPH_03516 |
| *flex6* | 3447 | T9SS C-terminal target domain-containing protein (REC74243.1) | KOACACPH_03517 |
| *flex7* | 1140 | EpsG family protein (SMC94123.1) | KOACACPH_03518 |
| *flex8* | 1011 | hypothetical protein | KOACACPH_03519 |
| *flex9* | 3654 | MMPL family transporter (WP_047492502.1) | KOACACPH_03520 |
| *flex10* | 915 | Dialkylrecorsinol condensing enzyme DarA (WP_123858105.1) | KOACACPH_03521 |
| *flex11* | 1140 | 3-oxoacyl-(acyl carrier protein) synthase III | KOACACPH_03522 |
| *flex12* | 432 | hypothetical protein | KOACACPH_03523 |
| *flex13* | 444 | ABC transporter permease (WP_090026350.1) | KOACACPH_03524 |
| *flex14* | 237 | hypothetical protein | KOACACPH_03525 |
| *flex15* | 2913 | Insulinase family protein (WP_084087117.1) | KOACACPH_03526 |
| *flex16* | 405 | hypothetical protein | KOACACPH_03527 |
| *flex17* | 996 | BtrH N-terminal domain-containing protein (WP_047492484.1) | KOACACPH_03528 |
| *flex18* | 759 | ABC transporter ATP-binding protein | KOACACPH_03529 |
| *flex19* | 1269 | ABC-2 type transporter | KOACACPH_03530 |
| *flex20* | 429 | Putative esterase | KOACACPH_03531 |
| *flex21* | 1134 | Beta-ketoacyl synthase | KOACACPH_03532 |
| *flex22* | 543 | 3-oxoacyl-ACP synthase (WP_081989140.1) | KOACACPH_03533 |
| *flex23* | 258 | Acyl carrier protein (WP_047492465.1) | KOACACPH_03534 |
| *flex24* | 1236 | Beta-ketoacyl synthase | KOACACPH_03535 |
| *flex25* | 1062 | Beta-ketoacyl synthase chain length factor (WP_047492459.1) | KOACACPH_03536 |
| *flex26* | 756 | Peptidoglycan-N-acetylglucosamine deacetylase | KOACACPH_03537 |
| *flex27* | 627 | Outer-membrane lipoprotein carrier protein | KOACACPH_03538 |
| *flex28* | 642 | hypothetical protein | KOACACPH_03539 |
| *flex29* | 597 | PorT family protein (WP_052188704.1) | KOACACPH_03540 |
| *flex30* | 381 | 3-hydroxyacyl-ACP dehydratase (WP_081989131.1) | KOACACPH_03541 |
| *flex31* | 477 | hypothetical protein | KOACACPH_03542 |
| *flex32* | 1170 | Undecaprenyl-phosphate 4-deoxy-4-formamido-L-arabinose transferase | KOACACPH_03543 |

| | | | |
|---|---|---|---|
| *flex33* | 2634 | T9SS-dependent M36 family metallopeptidase (WP_123321112.1) | KOACACPH_03544 |
| *flex34* | 2643 | T9SS-dependent M36 family metallopeptidase (WP_115919297.1) | KOACACPH_03545 |
| *flex35* | 366 | hypothetical protein | KOACACPH_03546 |
| *flex36* | 1686 | Acyl-CoA--6-aminopenicillanic acid acyl-transferase (WP_115919299.1) | KOACACPH_03547 |
| *flex37* | 1518 | NAD(P)/FAD-dependent oxidoreductase (WP_115919301.1) | KOACACPH_03548 |
| *flex38* | 885 | Lipid A biosynthesis acyltransferase (WP_047492440.1) | KOACACPH_03549 |
| *flex39* | 255 | Acyl carrier protein | KOACACPH_03550 |
| *flex40* | 1221 | Beta-ketoacyl synthase | KOACACPH_03551 |
| *flex41* | 732 | 3-oxoacyl-[acyl-carrier-protein] reductase FabG | KOACACPH_03552 |
| *flex42* | 1521 | Tyrosine 2,3-aminomutase | KOACACPH_03553 |
| *flex43* | 1257 | Tryptophan 7-halogenase (WP_084087087.1) | KOACACPH_03554 |
| *flex44* | 609 | hypothetical protein | KOACACPH_03555 |
| *flex45* | 1293 | Phenylacetate-coenzyme A ligase | KOACACPH_03556 |
| *flex46* | 1689 | M23 family metallopeptidase (WP_047492411.1) | KOACACPH_03557 |
| *flex47* | 3396 | Isoleucine--tRNA ligase | KOACACPH_03558 |
| *flex48* | 381 | RNA polymerase-binding transcription factor DksA | KOACACPH_03559 |
| *flex49* | 246 | Rhomboid family protein (WP_076356891.1) | KOACACPH_03560 |
| *flex50* | 195 | hypothetical protein | KOACACPH_03561 |
| *flex51* | 639 | Lipoprotein signal peptidase | KOACACPH_03562 |
| *flex52* | 645 | YdcF family protein (WP_047492396.1) | KOACACPH_03563 |
| *flex53* | 969 | Tryptophan--tRNA ligase | KOACACPH_03564 |
| *flex54* | 1284 | Sensor histidine kinase | KOACACPH_03565 |
| *flex55* | 672 | Phosphate regulon transcriptional regulatory protein PhoB | KOACACPH_03566 |
| *flex56* | 249 | hypothetical protein | KOACACPH_03567 |
| *flex57* | 450 | PepSY-like domain-containing protein (WP_034692507.1) | KOACACPH_03568 |
| *flex58* | 696 | Pyrimidine 5'-nucleotidase YjjG | KOACACPH_03569 |
| *flex59* | 135 | hypothetical protein | KOACACPH_03570 |
| *flex60* | 591 | RNA polymerase sigma-H factor | KOACACPH_03571 |
| *flex61* | 492 | hypothetical protein | KOACACPH_03572 |

**Table S2.** The description of microviridin biosynthesis related gene cluster from *Chryseobacterium* sp. KMC2

| Genes | Length (bp) | Functional annotation (Sequence ID from NCBI BLASTP) | Reference in genome |
|---|---|---|---|
| *mdnC* | 384 | Putative acyl-CoA thioester hydrolase | KOACACPH_03650 |
| *mdnD* | 798 | hypothetical protein | KOACACPH_03651 |
| *mdnE* | 2148 | Endonuclease MutS2 | KOACACPH_03652 |
| *mdnF* | 510 | GNAT family N-acetyltransferase (WP_084086267.1) | KOACACPH_03653 |
| *mdnG* | 642 | Uracil-DNA glycosylase | KOACACPH_03654 |
| *mdnH* | 504 | hypothetical protein | KOACACPH_03655 |
| *mdnI* | 543 | DUF456 domain-containing protein (WP_047487323.1) | KOACACPH_03656 |
| *mdnJ* | 1269 | Transporter, small conductance mechanosensitive | KOACACPH_03657 |
| *mdnK* | 720 | Pyridoxine 5'-phosphate synthase | KOACACPH_03658 |
| *mdnL* | 780 | Esterase YbfF | KOACACPH_03659 |
| *mdnA* | 240 | Microviridin (WP_084086262.1) | KOACACPH_03660 |
| *mdnB* | 219 | Microviridin (WP_047487341.1) | KOACACPH_03661 |
| *mdnM* | 957 | MvdC family ATP-grasp ribosomal peptide maturase (WP_084086261.1) | KOACACPH_03662 |
| *mdnN* | 1005 | MvdD family ATP-grasp ribosomal peptide maturase (WP_084086260.1) | KOACACPH_03663 |
| *mdnO* | 1014 | NAD-dependent epimerase/dehydratase family protein (WP_047487349.1) | KOACACPH_03664 |
| *mdnP* | 1770 | Long-chain-fatty-acid--CoA ligase FadD15 | KOACACPH_03665 |
| *mdnQ* | 1059 | Diphosphomevalonate decarboxylase (WP_084086257.1) | KOACACPH_03666 |
| *mdnR* | 942 | Endonuclease (WP_084086256.1) | KOACACPH_03667 |
| *mdnS* | 444 | Nuclear transport factor 2 family protein (WP_084086255.1) | KOACACPH_03668 |
| *mdnT* | 384 | VOC family protein (WP_047487892.1) | KOACACPH_03669 |
| *mdnU* | 384 | DUF1634 domain-containing protein (WP_047487366.1) | KOACACPH_03670 |
| *mdnV* | 831 | Sulfite exporter TauE/SafE family protein (WP_047487368.1) | KOACACPH_03671 |

**Table S3.** The description of siderophore biosynthesis related gene cluster from *Chryseobacterium* sp. KMC2

| Genes | Length (bp) | Functional annotation (Sequence ID from NCBI BLASTP) | Reference in genome |
|---|---|---|---|
| *speC* | 576 | Crp/Fnr family transcriptional regulator (WP_084084255.1) | KOACACPH_01784 |
| *speD* | 810 | Esterase family protein (WP_115917879.1) | KOACACPH_01785 |
| *speE* | 1098 | L-lysine 4-hydroxylase | KOACACPH_01786 |
| *speF* | 1512 | L-2,4-diaminobutyrate decarboxylase | KOACACPH_01787 |
| *speA* | 2409 | Putative siderophore biosynthesis protein | KOACACPH_01788 |
| *speH* | 1317 | L-lysine N6-monooxygenase | KOACACPH_01789 |
| *speB* | 1809 | IucA/IucC family siderophore biosynthesis protein (WP_047496558.1) | KOACACPH_01790 |
| *speI* | 1404 | MATE efflux family protein | KOACACPH_01791 |
| *speJ* | 246 | Acyl carrier protein (WP_084084248.1) | KOACACPH_01792 |
| *speK* | 1389 | 2-succinylbenzoate--CoA ligase | KOACACPH_01793 |

**Table S4.** The description of lanthipeptide I biosynthesis related gene cluster from *Chryseobacterium* sp. KMC2

| Genes | Length (bp) | Functional annotation (Sequence ID from NCBI BLASTP) | Reference in genome |
|---|---|---|---|
| *lan1* | 1191 | Alanine dehydrogenase | KOACACPH_00404 |
| *lan2* | 372 | hypothetical protein | KOACACPH_00405 |
| *lan3* | 774 | Histidine kinase (WP_084086149.1) | KOACACPH_00406 |
| *lan4* | 711 | Transcriptional regulatory protein NatR | KOACACPH_00407 |
| *lan5* | 378 | hypothetical protein | KOACACPH_00408 |
| *lan6* | 2289 | Sodium/hydrogen exchanger | KOACACPH_00409 |
| *lan7* | 654 | hypothetical protein | KOACACPH_00410 |
| *lan8* | 222 | Class I lanthipeptide (WP_047487795.1) | KOACACPH_00411 |
| *lan9* | 3036 | Lantibiotic dehydratase domain protein | KOACACPH_00412 |
| *lan10* | 1128 | Lanthionine synthetase C family protein | KOACACPH_00413 |
| *lan11* | 540 | hypothetical protein | KOACACPH_00414 |
| *lan12* | 417 | hypothetical protein | KOACACPH_00415 |
| *lan13* | 663 | hypothetical protein | KOACACPH_00416 |
| *lan14* | 843 | Cyclohexadienyl dehydrogenase | KOACACPH_00417 |
| *lan15* | 1419 | L-serine dehydratase 2 | KOACACPH_00418 |
| *lan16* | 579 | Transcription regulator, crp | KOACACPH_00419 |
| *lan17* | 624 | FMN-dependent NADH-azoreductase 1 | KOACACPH_00420 |
| *lan18* | 1344 | Ammonia channel | KOACACPH_00421 |
| *lan19* | 666 | Alpha/beta hydrolase (WP_115918317.1) | KOACACPH_00422 |
| *lan20* | 630 | Protein YceI | KOACACPH_00423 |
| *lan21* | 567 | Protein YceI | KOACACPH_00424 |

**Table S5.** The description of lanthipeptide II biosynthesis related gene cluster from *Chryseobacterium* sp. KMC2

| Genes | Length (bp) | Functional annotation (Sequence ID from NCBI BLASTP) | Reference in |
|---|---|---|---|
| *lan1* | 804 | HTH-type transcriptional activator RhaR | KOACACPH_02362 |
| *lan2* | 927 | Transcriptional regulator (WP_084085280.1) | KOACACPH_02363 |
| *lan3* | 636 | hypothetical protein | KOACACPH_02364 |
| *lan4* | 660 | hypothetical protein | KOACACPH_02365 |
| *lan5* | 600 | CDP-alcohol phosphatidyltransferase family protein (WP_115917644.1) | KOACACPH_02366 |
| *lan6* | 1449 | Asparagine--tRNA ligase | KOACACPH_02367 |
| *lan7* | 1464 | RNA polymerase sigma-54 factor | KOACACPH_02368 |
| *lan8* | 369 | hypothetical protein | KOACACPH_02369 |
| *lan9* | 1236 | hypothetical protein | KOACACPH_02370 |
| *lan10* | 3051 | Lantibiotic dehydratase domain protein | KOACACPH_02371 |
| *lan11* | 1158 | Lanthionine synthetase C family protein | KOACACPH_02372 |
| *lan12* | 918 | LLM class flavin-dependent oxidoreductase (WP_084085264.1) | KOACACPH_02373 |
| *lan13* | 186 | Class I lanthipeptide (WP_047488251.1) | KOACACPH_02374 |
| *lan14* | 891 | Carboxypeptidase-like regulatory domain-containing protein (WP_084085262.1) | KOACACPH_02375 |
| *lan15* | 1626 | 3-[(3aS,4S,7aS)-7a-methyl-1,5-dioxo-octahydro-1H-inden-4-yl]propanoyl:CoA | KOACACPH_02376 |
| *lan16* | 675 | Beta-carotene 15,15'-monooxygenase (WP_123942665.1) | KOACACPH_02377 |
| *lan17* | 492 | Sigma-70 family RNA polymerase sigma factor (WP_027374652.1) | KOACACPH_02378 |
| *lan18* | 531 | hypothetical protein | KOACACPH_02379 |
| *lan19* | 378 | hypothetical protein | KOACACPH_02380 |
| *lan20* | 870 | Polyphosphate:ADP phosphotransferase | KOACACPH_02381 |
| *lan21* | 465 | DUF1573 domain-containing protein (WP_047488266.1) | KOACACPH_02382 |

**Table S6.** The description of NRPS-like biosynthesis related gene cluster from *Chryseobacterium* sp. KMC2

| Genes | Length (bp) | Functional annotation (Sequence ID from NCBI BLASTP) | Reference in |
|---|---|---|---|
| *nrpsl1* | 1110 | DUF1624 domain-containing protein (WP_052188696.1) | KOACACPH_01527 |
| *nrpsl2* | 519 | Thioredoxin family protein (WP_047492101.1) | KOACACPH_01528 |
| *nrpsl3* | 396 | hypothetical protein | KOACACPH_01529 |
| *nrpsl4* | 357 | hypothetical protein | KOACACPH_01530 |
| *nrpsl5* | 144 | hypothetical protein | KOACACPH_01532 |
| *nrpsl6* | 1014 | Nucleoid-associated protein (WP_047492108.1) | KOACACPH_01533 |
| *nrpsl7* | 960 | hypothetical protein | KOACACPH_01534 |
| *nrpsl8* | 750 | Fumarate reductase iron-sulfur subunit | KOACACPH_01535 |
| *nrpsl9* | 1917 | Fumarate reductase flavoprotein subunit | KOACACPH_01536 |
| *nrpsl10* | 657 | Succinate dehydrogenase cytochrome b subunit (WP_047492118.1) | KOACACPH_01537 |
| *nrpsl11* | 1431 | Putative malate transporter YflS | KOACACPH_01538 |
| *nrpsl12* | 1164 | Porin (WP_084085123.1) | KOACACPH_01539 |
| *nrpsl13* | 1050 | Linear amide C-N hydrolase (WP_047492125.1) | KOACACPH_01540 |
| *nrpsl14* | 2193 | Sensor histidine kinase | KOACACPH_01541 |
| *nrpsl15* | 552 | Biliverdin-producing heme oxygenase (WP_047492131.1) | KOACACPH_01542 |
| *nrpsl16* | 927 | Malate dehydrogenase | KOACACPH_01543 |
| *nrpsl17* | 555 | hypothetical protein | KOACACPH_01544 |
| *nrpsl18* | 918 | Proline iminopeptidase | KOACACPH_01545 |
| *nrpsl19* | 3117 | Linear gramicidin synthase subunit D | KOACACPH_01546 |
| *nrpsl20* | 555 | Putative glycolipid-binding domain-containing protein (WP_157969889.1) | KOACACPH_01548 |
| *nrpsl21* | 573 | Crp/Fnr family transcriptional regulator (WP_047492147.1) | KOACACPH_01549 |
| *nrpsl22* | 537 | GNAT family N-acetyltransferase (WP_047492150.1) | KOACACPH_01550 |
| *nrpsl23* | 393 | AMP-dependent synthetase and ligase | KOACACPH_01551 |
| *nrpsl24* | 786 | hypothetical protein | KOACACPH_01552 |
| *nrpsl25* | 1647 | Alkaline phosphatase PafA | KOACACPH_01553 |
| *nrpsl26* | 1842 | Vitamin B12 transporter BtuB | KOACACPH_01554 |
| *nrpsl27* | 1059 | hypothetical protein | KOACACPH_01555 |
| *nrpsl28* | 1419 | CCA-adding enzyme | KOACACPH_01556 |
| *nrpsl29* | 348 | Nuclear transport factor 2 family protein (WP_047492172.1) | KOACACPH_01557 |
| *nrpsl30* | 549 | Threonylcarbamoyl-AMP synthase | KOACACPH_01558 |
| *nrpsl31* | 483 | IS200/IS605 family transposase (WP_115918710.1) | KOACACPH_01559 |
| *nrpsl32* | 513 | DinB family protein (WP_047492181.1) | KOACACPH_01560 |

| | | | |
|---|---|---|---|
| *nrpsl33* | 447 | GNAT family N-acetyltransferase (WP_047492184.1) | KOACACPH_01561 |
| *nrpsl34* | 1140 | Cystathionine beta-lyase | KOACACPH_01562 |
| *nrpsl35* | 324 | Gliding motility protein GldC (WP_047492191.1) | KOACACPH_01563 |
| *nrpsl36* | 984 | Gliding motility protein GldB (WP_081989115.1) | KOACACPH_01564 |
| *nrpsl37* | 639 | Internalin J | KOACACPH_01565 |
| *nrpsl38* | 522 | Acetyltransferase | KOACACPH_01566 |
| *nrpsl39* | 531 | N-acetyltransferase (WP_115918805.1) | KOACACPH_01567 |
| *nrpsl40* | 792 | NH(3)-dependent NAD(+) synthetase | KOACACPH_01568 |
| *nrpsl41* | 516 | Ribonuclease | KOACACPH_01569 |

**Table S7.** The description of genes and metabolic pathways related to keratin utilization in *Chryseobacterium* sp. KMC2 genome

| Gene order | EC number | KO number | Functional annotation | Metabolic pathway | Reference in genome |
|---|---|---|---|---|---|
| 1 | 2.6.1.42 | K00826 | Branched-chain-amino-acid aminotransferase | Metabolism of Leucine and Isoleucine | KOACACPH_00391/KOACACPH_04122 |
| 2 | 1.2.4.4 | K11381 | 3-methyl-2-oxobutanoate dehydrogenase | Metabolism of Leucine and Isoleucine | KOACACPH_00351 |
| 3 | 2.3.1.168 | K09699 | Dihydrolipoyllysine-residue (2-methylpropanoyl)transferase | Metabolism of Leucine and Isoleucine | KOACACPH_03134 |
| 4 | 1.3.8.7 | K00248, K00249 | Acyl-CoA dehydrogenase | Metabolism of Leucine and Isoleucine | KOACACPH_01887 |
| 5 | 4.2.1.18 | K13766 | Methylglutaconyl-CoA hydratase | Metabolism of Leucine and Isoleucine | KOACACPH_03720 |
| 6 | 4.1.3.4 | K01640 | Hydroxymethylglutaryl-CoA lyase | Metabolism of Leucine and Isoleucine | KOACACPH_03960 |
| 7 | 1.3.8.1 | K00248 | Acyl-CoA dehydrogenase | Metabolism of Leucine and Isoleucine | KOACACPH_00342/KOACACPH_03381 |
| 8 | 4.2.1.17 | K01715, K01692 | Enoyl-CoA hydratase | Metabolism of Leucine and Isoleucine | KOACACPH_03576/KOACACPH_01888/KOACACPH_01890 |
| 9 | 1.1.1.35 | K01782, K07516 | 3-hydroxyacyl-CoA dehydrogenase | Metabolism of Leucine and Isoleucine | KOACACPH_03378 |
| 10 | 2.3.1.16 | K00623 | Acetyl-CoA acyltransferase | Metabolism of Leucine and Isoleucine | KOACACPH_03380 |
| 11 | 2.3.3.1 | K01647 | Citrate synthase | TCA cycle | KOACACPH_01094/KOACACPH_01792 |
| 12 | 4.2.1.3 | K01681 | Aconitase | TCA cycle | KOACACPH_03590 |
| 13 | 1.1.1.42 | K00031 | Isocitrate dehydrogenase | TCA cycle | KOACACPH_00544 |
| 14 | 1.2.4.2 | K00164, K01616 | Oxoglutarate dehydrogenase | TCA cycle | KOACACPH_00825 |
| 15 | 2.3.1.61 | K00658 | Dihydrolipoyl succinyltransferase | TCA cycle | KOACACPH_00826 |
| 16 | 1.8.1.4 | K00382 | Dihydrolipoyl dehydrogenase | TCA cycle | KOACACPH_00132/KOACACPH_00765 |
| 17 | 6.2.1.5 | K01902, K01903, K02381 | Succinate-CoA ligase | TCA cycle | KOACACPH_01458/KOACACPH_03423 |
| 18 | 2.8.3.18 | K01067, K18118 | Succinyl-CoA:acetate CoA-transferase | TCA cycle | KOACACPH_02425 |
| 19 | 1.3.5.1 | K00240 | Succinate dehydrogenase | TCA cycle | KOACACPH_01535/KOACACPH_01536/KOACAC |

| | | | | | |
|---|---|---|---|---|---|
| | | | | | PH_02931/KOACACPH_02932 |
| 20 | 4.2.1.2 | K01676, K01677, K01678, K01679 | Fumarate hydratase | TCA cycle | KOACACPH_02695/KOACACPH_02696 |
| 21 | 1.1.5.4 | K00116 | Malate dehydrogenase (quinone) | TCA cycle | KOACACPH_02576 |
| 22 | 1.1.1.37 | K00024 | Malate dehydrogenase | TCA cycle | KOACACPH_01543 |
| 23 | 4.1.1.49 | K01610 | Phosphoenolpyruvate carboxykinase (ATP) | Glycolysis/Gluconeogenesis | KOACACPH_00978 |
| 24 | 4.2.1.11 | K01689 | Phosphopyruvate hydratase | Glycolysis/Gluconeogenesis | KOACACPH_01093 |
| 25 | 5.4.2.11 | K01834 | 2,3-bisphosphoglycerate-dependent phosphoglycerate mutase | Glycolysis/Gluconeogenesis | KOACACPH_01759 |
| 26 | 5.4.2.12 | K15633 | 2,3-bisphosphoglycerate-independent phosphoglycerate mutase | Glycolysis/Gluconeogenesis | KOACACPH_03235 |
| 27 | 2.7.2.3 | K00927 | Phosphoglycerate kinase | Glycolysis/Gluconeogenesis | KOACACPH_03849 |
| 28 | 1.2.1.12 | K00134 | Glyceraldehyde-3-phosphate dehydrogenase | Glycolysis/Gluconeogenesis | KOACACPH_03436 |
| 29 | 1.2.1.9 | K00131 | NADP-dependent glyceraldehyde-3-phosphate dehydrogenase | Glycolysis/Gluconeogenesis | KOACACPH_00524 |
| 30 | 4.1.2.13 | K01624 | Fructose-bisphosphate aldolase | Glycolysis/Gluconeogenesis | KOACACPH_03902/KOACACPH_04210 |
| 31 | 2.7.1.11 | K00850, K21071 | 6-phosphofructokinase | Glycolysis/Gluconeogenesis | KOACACPH_03435 |
| 32 | 3.1.3.11 | K03841 | Fructose-bisphosphatase | Glycolysis/Gluconeogenesis | KOACACPH_01662 |
| 33 | 5.4.2.2 | K01835 | Phosphoglucomutase | Glycolysis/Gluconeogenesis | KOACACPH_03206 |
| 34 | 2.7.1.40 | K00873 | Pyruvate kinase | Pyruvate metabolism | KOACACPH_02949 |
| 35 | 1.2.4.1 | K00161, K00162 | pyruvate dehydrogenase | Pyruvate metabolism | KOACACPH_03262/KOACACPH_02307 |
| 36 | 2.3.3.5 | K01659 | 2-methylcitrate synthase | Propanoate metabolism | KOACACPH_01884 |
| 37 | 4.2.1.79 | K01720 | 2-methylcitrate dehydratase | Propanoate metabolism | KOACACPH_01885 |
| 38 | 4.2.1.99 | K01682 | 2-methylaconitate hydratase | Propanoate metabolism | KOACACPH_03589 |
| 39 | 1.2.4.4 | K11381 | 3-methyl-2-oxobutanoate dehydrogenase | Propanoate metabolism | KOACACPH_00351 |
| 40 | 2.3.1.168 | K09699 | Dihydrolipoyllysine-residue (2-methylpropanoyl)transferase | Propanoate metabolism | KOACACPH_03134 |
| 41 | 4.1.3.30 | K03417 | 2-methylisocitrate lyase | Propanoate metabolism | KOACACPH_01883 |

| | | | | | |
|---|---|---|---|---|---|
| 42 | 2.5.1.6 | K00789 | S-adenosylmethionine synthase | Metabolism of Methionine and Cysteine | KOACACPH_03573 |
| 43 | 2.1.1.37 | K00558 | DNA (cytosine-5-)-methyltransferase | Metabolism of Methionine and Cysteine | KOACACPH_02976 |
| 44 | 3.3.1.1 | K01251 | Adenosylhomocysteinase | Metabolism of Methionine and Cysteine | KOACACPH_03683 |
| 45 | 4.2.1.22 | K01697 | Putative cystathionine beta-synthase | Metabolism of Methionine and Cysteine | KOACACPH_01627 |
| 46 | 4.4.1.1 | K01758 | L-methionine gamma-lyase | Metabolism of Methionine and Cysteine | KOACACPH_01562 |
| 47 | 2.6.1.1 | K00812 | Aspartate/prephenate aminotransferase | Metabolism of Phenylalanine, Tyrosine, Glutamate and Aspartate | KOACACPH_02216 |
| 48 | 4.3.1.1 | K01744 | Aspartate ammonia-lyase | Metabolism of Phenylalanine, Tyrosine, Glutamate and Aspartate | KOACACPH_04295 |
| 49 | 4.1.1.19 | K01585 | Biosynthetic arginine decarboxylase | Metabolism of Arginine | KOACACPH_01491 |
| 50 | 3.5.3.11 | K01480 | Agmatinase | Metabolism of Arginine | KOACACPH_01493 |
| 51 | 2.5.1.16 | K00797 | Polyamine aminopropyltransferase | Metabolism of Arginine | KOACACPH_01104 |
| 52 | 1.5.99.6 | K00316 | Spermidine dehydrogenase | Metabolism of Arginine | KOACACPH_01105 |
| 53 | 1.2.1.3 | K00128 | Aldehyde dehydrogenase (NAD+) | Metabolism of Arginine | KOACACPH_01832/KOACACPH_01990 |
| 54 | 2.6.1.19 | K07250,K13524 | 4-aminobutyrate aminotransferase | Metabolism of Arginine | KOACACPH_03346 |
| 55 | 1.2.1.24 | K08324 | Succinate-semialdehyde dehydrogenase (NAD+) | Metabolism of Arginine | KOACACPH_02948 |
| 56 | 1.2.1.79 | K00135 | Succinate-semialdehyde dehydrogenase (NADP+) | Metabolism of Arginine | KOACACPH_02948 |
| 57 | 2.7.7.4 | K00955 | Sulfate adenylyltransferase | Sulfur metabolism | KOACACPH_01861/KOACACPH_01862 |
| 58 | 2.7.1.25 | K00955 | Adenylyl-sulfate kinase | Sulfur metabolism | KOACACPH_01862 |
| 59 | 1.8.4.8 | K00390 | Phosphoadenosine phosphosulfate reductase | Sulfur metabolism | KOACACPH_01860 |
| 60 | 1.8.1.2 | K00380 | Sulfite reductase (NADPH) flavoprotein alpha-component | Sulfur metabolism | KOACACPH_01866 |
| 61 | 1.8.1.2 | K00381 | Sulfite reductase (NADPH) hemoprotein beta-component | Sulfur metabolism | KOACACPH_01867 |
| 62 | 1.8.7.1 | K00392 | Assimilatory sulfite reductase | Sulfur metabolism | KOACACPH_01867 |
